# Enhanced deviant responses in patterned relative to random sound sequences

**DOI:** 10.1101/214031

**Authors:** Rosy Southwell, Maria Chait

## Abstract

How are brain responses to deviant events affected by the statistics of the preceding context? We recorded electroencephalography (EEG) brain responses to frequency deviants in matched, regularly-patterned (REG) versus random (RAND) tone-pip sequences. Listeners were naïve and distracted by an incidental visual task. Stimuli were very rapid so as to limit conscious reasoning about the sequence order and tap automatic processing of regularity.

Deviants within REG sequences evoked a substantially larger response (by 71%) than matched deviants in RAND sequences from 80 ms after deviant onset. This effect was underpinned by distinct sources in right temporal pole and orbitofrontal cortex in addition to the standard bilateral temporal and right pre-frontal network for generic frequency deviance-detection. These findings demonstrate that the human brain rapidly acquires a detailed representation of regularities within the sensory input and evaluates incoming information according to the context established by the specific pattern.

## Introduction

Detection of new events within a constantly fluctuating sensory input is a fundamental challenge to organisms in dynamic environments. Hypothesized to underlie this process is a continually-refined internal model of the real-world causes of sensations, made possible by exploiting statistical structure in the sensory input^1-4^. Evidence from multiple domains, including speech^5^, abstract sound sequences^6,7^, vision^8^ and motor control^9^ reveals sensitivity to environmental statistics which influences top-down, expectation-driven perceptual processing. When the organism encounters a sensory input that is inconsistent with the established internal model, and is therefore indicative of a potentially relevant change in the environment, a ‘prediction error’ or ‘surprise’ response is generated^10^, promoting a rapid reaction to the associated environmental change. Understanding what aspects of stimuli are ‘surprising’, is therefore central to understanding this network.

The auditory system has been a fertile ground for probing sensory error responses, at multiple levels of the processing hierarchy^11-13^. A common approach involves using a stream of standard sounds to establish a regularity that is occasionally interrupted by ‘deviant’ sounds^14-17^. Deviants usually evoke an increased response relative to that measured for the standards^16,18,19^. Since many of the investigated sequences have been very simple, often a repeated tone, neural adaptation is likely a major contributor to this process^12,20,21^. However, accumulating evidence suggests that at least part of the response arises from neural processes associated with computing ‘surprise’ or detecting a mismatch between expected and actual sensory input^15,22,23^

What is the information used in calculating surprise? By modelling brain responses to two-tone sequences of different base probabilities, Rubin et al.^4^ demonstrated that trial-wise neural responses in auditory cortex are well explained by the probability of occurrence of each tone frequency, calculated from the recent history of the sequence. The models that best fit neural responses were based on a relatively long stimulus history (~10 tones) but a coarse representation. In line with this conclusion, Garrido et al.^24^ demonstrated that MEG responses to probe tones are sensitive to the statistical context (mean and variance of frequency) of randomly generated tone-pip sequences such that larger responses occurred to the same probe tone when presented in a context with low, as opposed to high, frequency variance. Similarly, Daikhin & Ahissar^22^ measured responses to frequency deviants embedded into a sequence of fixed standards or standards that were slightly jittered in frequency. Whereas the response to the standard was not affected by this manipulation, deviant responses were reduced in the jittered condition. In both cases^22,24^ the results were interpreted as the deviant response showing sensitivity to the uncertainty induced by the sensory context, such that deviants within a more volatile (less predictable) context are considered less surprising than identical events within a more stable, or precise background (see also Hermann et al.^19^).

Overall, mounting evidence suggests that the deviant response is shaped by the dynamic statistics of the unfolding sequence. However, relatively little is known about the properties of this representation of the past: i.e. whether it is coarse, reflecting a small set of summary statics (e.g. as suggested by Rubin et al^4^), and possibly underpinned by adaptation processes^15,19,25^, or instead keeping track of a more detailed history.

Here we investigated whether error responses to frequency deviants, within complex sound sequences, are affected by the specific temporal patterning of the preceding sequence. We used rapid tone-pip sequences, unique on each trial, that occasionally contained a frequency deviant presented outside of the frequency region occupied by the standards. To understand whether the deviant response merely reflects an unexpected change in frequency between the standards and deviant, or whether it is also affected by the temporal *order* of elements in the sequence, we used either regular (REG) or random (RAND) sequences of otherwise matched frequencies (see Figure 1). The precision of the available information regarding successive frequencies can be either low (RAND) or high (REG). Importantly, overall frequency occurrence statistics, taken over the sequence duration or over the entire experimental session, are identical between REG and RAND. The resulting effect is that the context offered by each sequence differs in predictability but not in frequency span.

**Figure 1:**
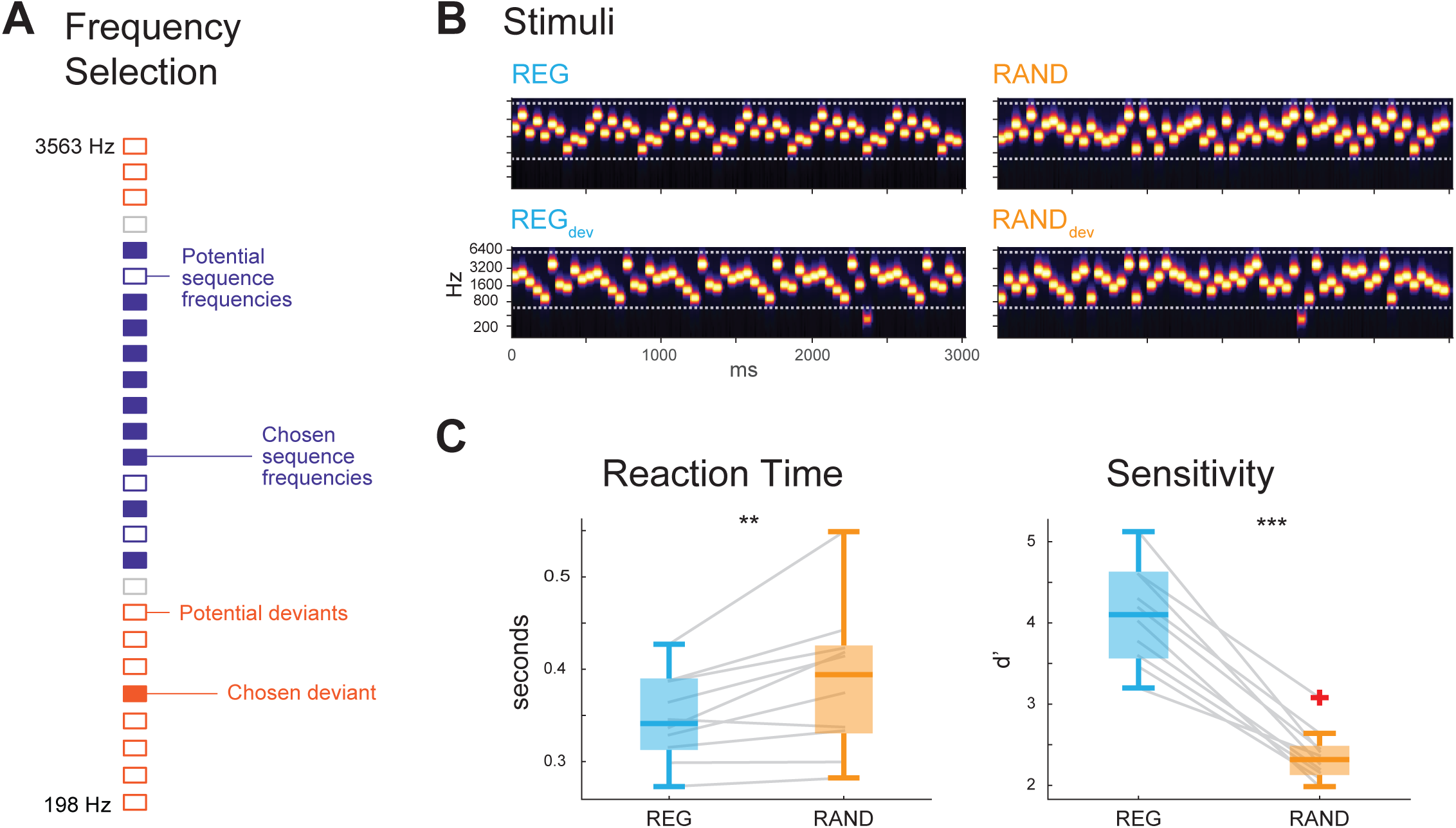
Stimuli and Behavioural responses. **a:** Procedure for selecting frequencies used for each stimulus. From the pool of 26; 13 adjacent values were chosen at random as candidate sequence frequencies (purple); 10 were selected for the sequence. Of the remaining tones; all except the frequencies closest to the sequence could potentially be deviants (orange); and from these a single value was chosen at random to be the deviant on that trial. **b:** Example set of stimuli for the four conditions; these were generated together from the same frequencies in order to match acoustic properties. **c:** Results from the behavioural experiment. **Left:** reaction times to deviant tones. **Right:** sensitivity (d’) to deviant tones. ** p < 0.01, ***p < 0.001

We find that deviant tones are detected more effectively in REG than in RAND sequences, as revealed both behaviourally and by measuring deviant-evoked EEG responses in naïve, distracted listeners. These results confirm that the human brain tracks and evaluates incoming sensory information against the specific pattern established by the sequence context. Notably, this occurs even within very rapid sound sequences where conscious reasoning about the sequence order is unlikely to be possible.

## Results

### Experiment 1 - Behavioural sensitivity to deviants in REG and RAND sequences

We measured listeners’ ability to detect frequency deviants in matched REG and RAND sequences (Figure 1a; b). The mean reaction time to deviants in a control condition (CTRL), where all tone-pips except the deviant were at the same frequency, was 329±16 ms, giving an estimate of participants’ basic response time. The mean reaction times to REG and RAND deviants were 347±15 ms and 387±25 ms, respectively. Paired-sample t-tests were carried out on the subject-wise averages of both RT and d’ for REG versus RAND. Reaction times were significantly faster (p = 0.01) and sensitivity (d’) significantly higher (p < 0.001) to deviants in REG, versus RAND sequences. See Figure 1c.

To summarise, despite carefully matched properties of the regular and random stimuli used, we observe robustly greater behavioural sensitivity, as well as faster reaction times, to deviant tones which violate a regular sequence.

### Experiment 2 - EEG in naïve, passively listening participants

EEG responses were recorded to sequences identical to those used in Experiment 1 (REG, RAND, REG_dev_ and RAND_dev_). In order to capture automatic, stimulus-driven deviance detection processes, participants were kept naïve and distracted, watching a silent, subtitled movie of their choice.

### Post-session reports

Following the EEG experiment, participants were questioned about the sounds presented (see Methods). Nine out of twenty described hearing some kind of pattern in the sound, for instance ‘repetition’ and ‘alternating high and low sounds’, although these descriptions were usually quite vague, and when pressed to elaborate, none had noticed the distinction between REG and RAND trials. Thirteen subjects reported hearing occasional sounds which broke the pattern, or were otherwise distinctive; and when asked to elaborate, several specified that the pitch of the tones stood out as higher or lower than the rest. This shows that the deviants entered subjects’ awareness at least in some cases, although accurate description of the patterning of the sequences was much rarer. The mean rating given for how distracting the sound sequences were overall was 2.2 out of 5, range 1-4; indicating that subjects were moderately distracted by the sound sequences on average, but with considerable variability across the group.

### EEG

#### Deviant-evoked responses

The deviant-evoked responses (Figure 2a) were comprised of a series of peaks closely resembling the standard N1-P2-N2 sequence commonly observed at stimulus onset, or for changes within ongoing sounds^26^. To statistically assess these effects, firstly channels and time-intervals showing a significant effect of deviance were identified with the contrast (REG_dev_ + RAND_dev_) – (REG + RAND), thus defining a region-of-interest (ROI) in time-channel space (see ‘Methods’). This allowed separation of neural activity associated with the ongoing context of the sequence from those strictly evoked by the deviant. The resulting three clusters, shown in Figure 2b, correspond to the peaks observed in the time domain (Figure 2a). The first cluster was a fronto-central negativity between 80 and 145 ms (p = 0.001), corresponding most closely in time and topography to the N1. The second cluster at 165-245 ms (p = 0.001) had a similar topography but reversed polarity, and a third cluster from 290 to 320 ms (p = 0.016) had a smaller spatial extent and negative polarity (Figure 2c).

**Figure 2:**
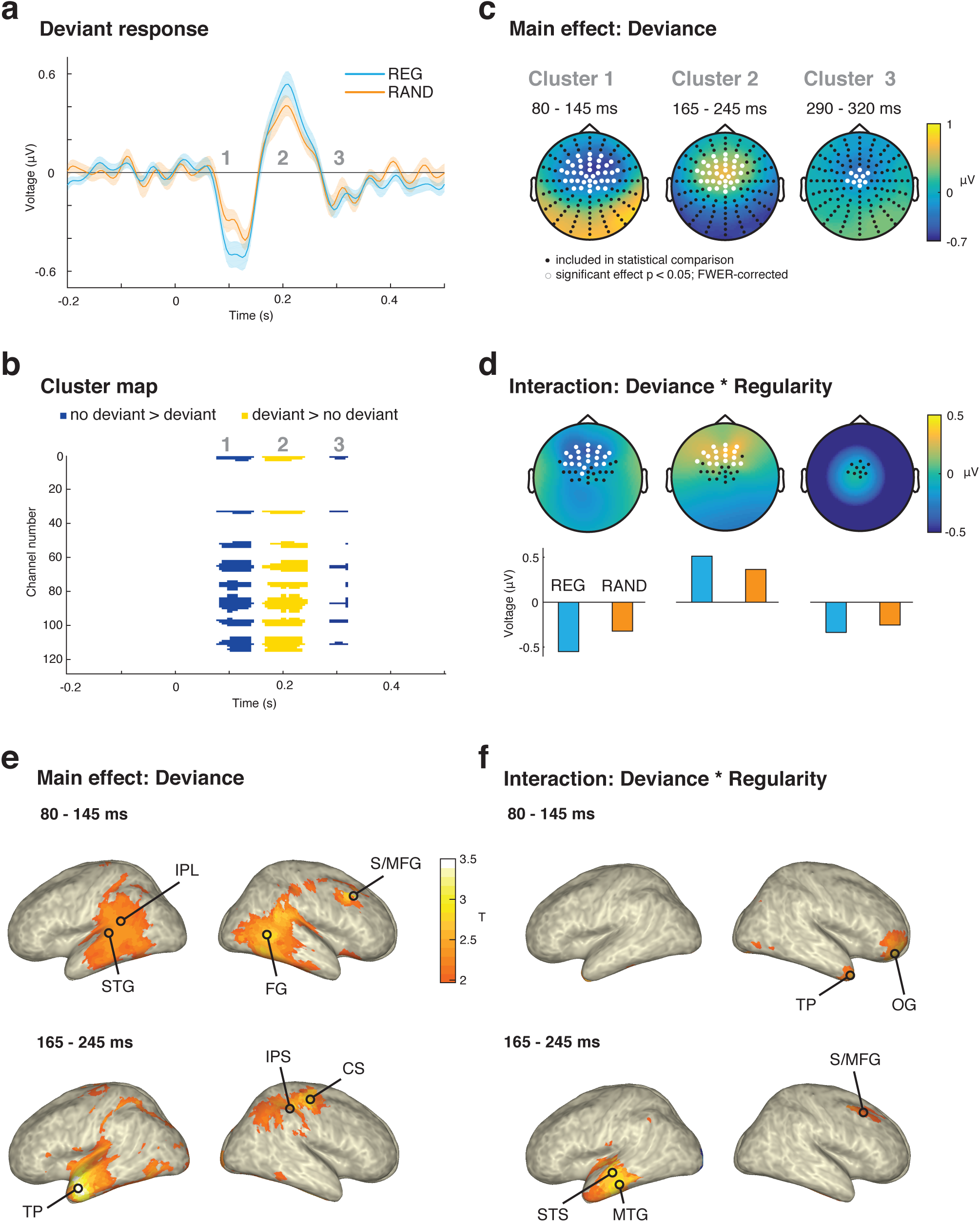
Deviant-evoked response. **a:** Time-domain response averaged over 39 central channels which show a main effect of deviance. Error bars show standard error in the mean over subjects computed by bootstrap resampling. Three deflections in the response correspond to the three clusters shown in (b). **b:** three clusters showing a main effect of deviance; i.e. (REG_dev_ + RAND_dev_) – (REG + RAND) computed over all time-points and channels shown. **c:** Topography of the three main-effect clusters; averaged over the temporal extent of each cluster. **d:** topography of the interaction contrast (REG_dev_ – REG) – (RAND_dev_ – RAND). Channels included in the statistical analysis are shown in black (these are the significant channels in (c)). Channels showing an effect of deviance at any point during the cluster are highlighted in white. The average magnitude of the response to REG_dev_ and RAND_dev_, within the ROI defined by the main effect cluster, is shown in the bar plots below. **e,f:** Source-level activity shown on a template cortical sheet. T-statistic maps thresholded at T = 2. All show average source activity taken over a time-window defined by significant effect of deviance in sensor space. **e:** Effect of deviance from 80-145 ms (**top**) and from 165-245 ms (**bottom**). **f:** effect of regularity on the deviant response from 80-145 ms (**top**) and from 165-245 ms (**bottom**).

The orthogonal contrast of the deviant response magnitude by sequence type, i.e. (REG_dev_ – REG) – (RAND_dev_ – RAND) was then calculated for the ROI defined by the time points and channels in each of the three clusters identified above. Statistical analysis was performed using a cluster-based permutation test to correct for multiple comparisons (see Methods). In the first cluster, a subset (21 channels) of the ROI showed an effect of regularity on the deviant response (p = 0.005), which was 71% larger (calculated over mean activity within the significant channels), in REG sequences. The second cluster, similar to a P2 component, was also larger (by 41%) in REG (p = 0.002) in a subset of 17 channels (Figure 2d). There was no effect of regularity on the third cluster. Importantly, since the analysis above is performed on high pass filtered, and baselined, data, the effect of regularity on the deviance response occurs over and above the sustained response difference between the two sequence types (see below).

For the main effect of deviance, extensive activity in the first time window was observed in bilateral temporal cortex, including auditory areas within superior temporal gyrus (STG) and spreading to parietal cortex (Figure 2e, top). Additionally, right superior/middle frontal gyrus (S/MFG) showed a deviant-evoked response, containing the maximal T-statistic of 3.05 (Figure 2e, top). During the second window, the deviant response was associated with temporal lobe activation, but this time more prominently left-lateralised as well as situated more frontally around the temporal pole (TP), with a peak of T = 3.55 in the left middle temporal gyrus. Right-hemisphere activation is seen around the intraparietal sulcus (IPS) and the central sulcus (CS; Figure 2e, bottom).

For the interaction of the deviant response with regularity from 80-145 ms (Figure 2f, top), we observed increased activity to REG deviants at right TP and right orbital gyrus (OG), where the maximal t-statistic of 2.86 was observed. From 165-245 ms, REG_dev_ elicited greater activity than RAND_dev_ in left temporal cortex, with a peak T-statistic of 3.05 in left middle temporal gyrus (MTG) and superior temporal sulcus (STS). Increased activity was also seen in right S/MFG.

#### Sequence Offset responses

Interestingly, an effect of regularity is also present during the offset response, which is seen from about 50 ms after the cessation of the sequence (Figure 3a). The offset peak was compared between REG and RAND (collapsed across deviant and no-deviant trials) using the same clustering approach as above. REG showed a significantly larger offset response than RAND, from 85-175 ms (p < 0.001) in most channels (more negative in a fronto-central cluster of 58 channels, p < 0.001; and more positive in a temporal-occipital cluster of 50 channels, p < 0.001). There was also a significantly more positive response from 215-300 ms (p = 0.008) post-offset in a fronto-central cluster of 41 channels (Figure 3b; lower right).

**Figure 3:**
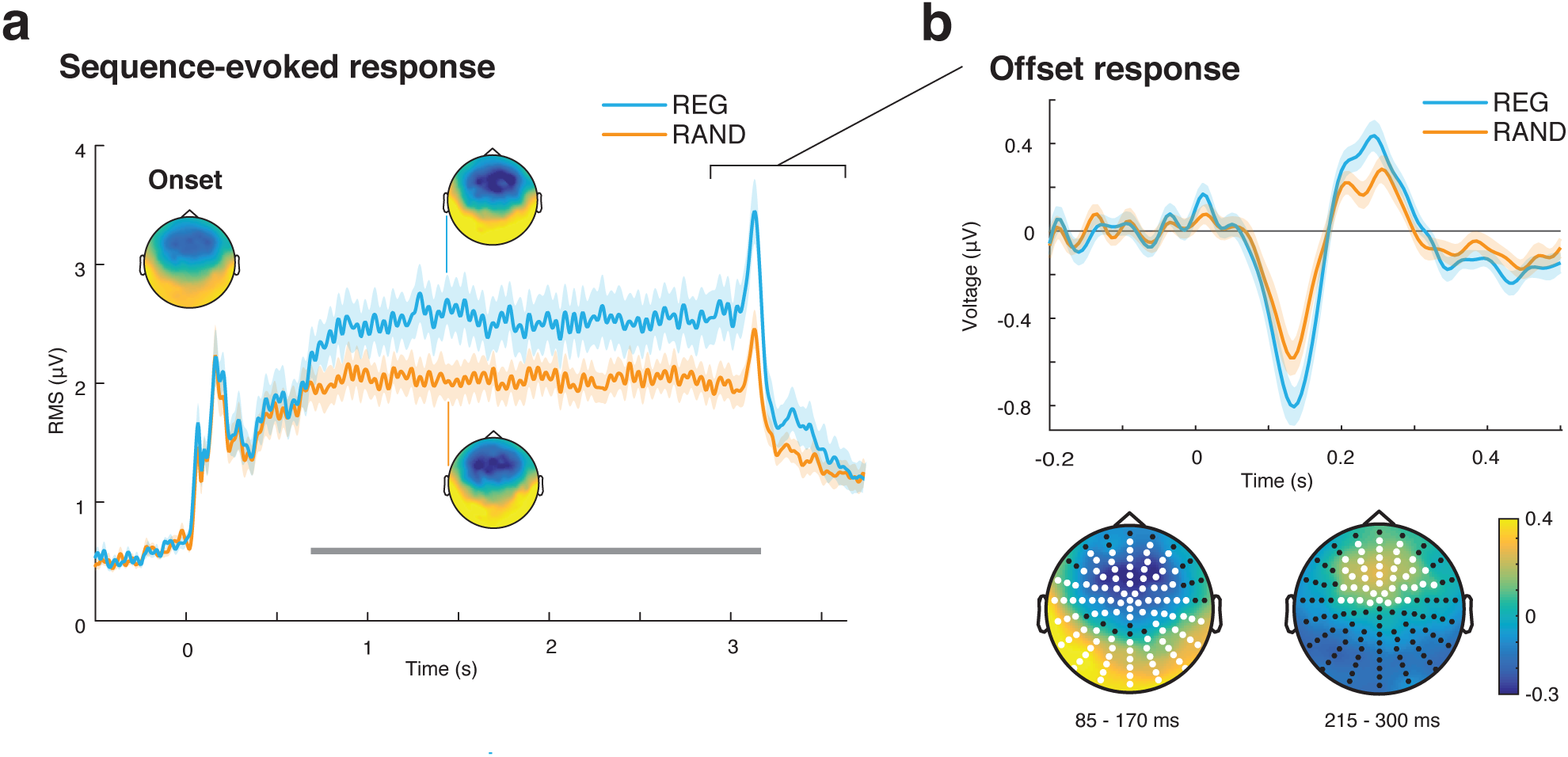
Sequence-evoked responses. **a.** Sequence-evoked response. Shown is the root-mean-square (RMS) over all channels; error bars show standard error of the mean over subjects computed by bootstrap resampling. Time period showing significant difference between REG and RAND conditions is indicated by a grey bar. Topography of the onset response is shown (inset; left) for comparison to the topography of the sustained response to REG (inset; top) and RAND (inset; bottom). **b:** Offset response. **Top:** Evoked response averaged over 58 central channels showing an effect of regularity. **Bottom:** Topography of the response during the two time-windows covering significant clusters for the contrast (REG – RAND); channels showing an effect of deviance at any point during the cluster are highlighted in white.

Statistical comparison was performed at each time-point and channel, but for illustrative purposes the time-domain response averaged over the 58 channels in the first negative cluster, is shown in Figure 3b.

#### Sequence evoked responses

The sequence-evoked response is shown in Figure 3a; deviant and no-deviant conditions were pooled and REG and RAND conditions are plotted separately. The standard sequence of auditory onset responses is seen, followed by a rise to a sustained response that persists until stimulus offset. The topography of this response for both REG and RAND is similar to the N1 onset response, namely a fronto-central negativity (see inset topographies; Figure 2c). The response to REG was significantly greater than that to RAND, from 705 ms after onset until 440 ms after offset (p < 0.001, FWER-corrected). The response to REG diverged from RAND after just 4 tone-pips (200 ms) of the first repeated cycle. This pattern of results entirely replicates previous work^27,28^ and is consistent with ideal-observer-like sensitivity to the emergence of regularity^27^. However, the present stimuli are better controlled for effects of frequency-specific adaptation, by ensuring that REG and RAND have exactly the same frequency content; and by disallowing repetitions of the same frequency on two adjacent tone-pips.

## Discussion

We investigated whether the deterministic predictability of successive events within rapid tone-pip sequences influences responses to deviant tones. Whilst the fact that regularity shapes responses to standards is a commonly observed effect, even in complex sequences^14,15^, effects on the deviant response have been more elusive. For example, Yaron et al.^29^ report remarkable sensitivity to the temporal patterning of long sound sequences, but these effects are revealed via changes to the response to the standard, but not deviant sounds. Similarly, Costa-Faidella et al.^30^ showed robust effects of regularity on the standard, such that more repetition suppression is seen in a temporally regular than a jittered context - but the response to the deviant itself did not differ (see also Christianson et al.^31^). In contrast, we demonstrate that early (from 80 ms) EEG responses in naïve, distracted listeners are significantly affected by sequence regularity such that deviants within regular sequences evoked a larger response than matched deviants in random sequences. The effect of regularity was also revealed behaviourally - listeners are faster and substantially more accurate at detecting deviant events within regularly repeating (REG), relative to random (RAND) tone-pip sequences, despite matched frequency content.

Increased responses to deviants within complex acoustic patterns have previously been reported in the MMN^32^, statistical learning and music processing literature. For example, the ERAN (Early right anterior negativity) is a brain response, seen from around 150 ms following chord onset, commonly evoked by music syntax irregularities^33,34^. Further, in a study of musical expectation, Pearce et al.^35^ showed that low probability notes, compared to high probability notes, elicited a larger negative component at around 400 ms. Using non-musical, abstract sound sequences with specifically controlled transition probabilities, Koelsch et al.^36^ showed increased negativity to less probable items from 130-220 ms after onset. Furl et al.^37^ trained participants to discriminate Markov sequences of pure tones from random ones and demonstrated a difference between low and high probability tones from 200 ms post-onset (during the P2 peak) originating in the right temporo-parietal junction.

However, this previous work has used regularities established over an extended period (e.g. a fixed transition probability matrix throughout the experiment; or even – for music - over a lifetime) and brain activity was often recorded while participants were consciously making decisions about the predictability of the pattern^35,37^, therefore potentially involving longer-term learning mechanisms supported by a different network than the rapid, automatic and pre-attentive processes studied here. Indeed, the latencies reported from such work tend to be later than those observed here.

The deviant responses seen here - a standard succession of N1-P2-N2 deflections - are similar to those commonly observed in the human Stimulus-Specific Adaptation (SSA) literature^21,38^ and which have previously been shown to be affected by both simple adaptation (repetition suppression) as well as more complex statistical context (relative probability of the deviant). Here we demonstrate a substantially larger deviant response (71% increase in the first window) in REG relative to RAND sequences, confirming that these early deviant-evoked responses are also subject to automatic modulation by the degree of predictability in the ongoing sequence context. These findings are consistent with the notion that the brain continually tracks and maintains a detailed representation of the structure of the unfolding sensory input and that this representation shapes the processing of incoming information.

An alternative explanation for the observed findings might have been that regular patterns automatically attract attention^39^, and that this facilitates the detection of deviants in REG sequences. Southwell et al.^28^ directly investigated the question of whether attention is biased towards REG sequences (essentially identical to those used here), and found no attentional bias towards either REG or RAND. The fact that when interrogated, participants in the present study did not report noticing a distinction between REG and RAND trials also supports the conclusion that attention is not a likely explanation for the observed pattern of effects. Furthermore, the effects of attention on deviance detection are commonly associated with the presence of a P300 response^40,41^ reflecting the fact that the deviant was consciously perceived. The P300 was absent here. Instead our results point to an early and time-limited (between 80-250ms) effect of context on the deviant response.

We also observed a remarkably strong effect of regularity on the offset response to the sequences. An offset is a special case of deviance, reflecting the violation of the expectation that a tone will be presented. This effect has been studied extensively in the context of the auditory omission^42,43^ or offset^44^ paradigms, where an evoked response occurs to unexpected omissions of sounds, at a similar latency to the early responses to actual sounds, but only when the preceding sequence allowed a prediction to be formed about the omitted tone’s properties. That both frequency and offset deviants are affected by regularity is consistent with the notion that the overall predictability of the pattern (the precision of the prediction the observer can make about an upcoming event) affects error responses regardless of the dimension in which the deviance occurs.

The main effect of deviance, collapsed over sequence context and hence assumed to reflect the mismatch in frequency, was significant across a central subset of channels commonly associated with auditory responses (Figure 2c). Source analysis revealed that activity within the early time window (80-145 ms) originated in temporal cortex and right prefrontal cortex. This is consistent with the standard network of bilateral auditory and right-hemisphere frontal sources often implicated in pre-attentive deviance detection^16,45-47^. In the later time window (165-245 ms), the anterior portion of the left temporal cortex showed the strongest deviant-evoked response, with some additional activation in right intraparietal sulcus (IPS). The IPS is commonly implicated in auditory perceptual organisation^48^ and specifically figure-ground segregation^49^ and its involvement here may be linked to processes which stream the deviant tone away from the ongoing sequence.

The increased response to deviants in REG sequences was associated with regions that are, at least in part, distinct from those involved in coding for the frequency mismatch (main effect of deviance). This is consistent with the observation that the deviance-by-regularity effect was only significantly present in a frontal subset of the channels activated by the main effect of deviance. Notably, whereas the main effect of deviance in the early time window was dominated by extensive activation of temporal areas, the effect of sequence regularity is mostly seen in the right temporal pole and right orbitofrontal cortex. Both of the latter have previously been implicated in sensitivity to context: the right anterior temporal cortex has been shown to be sensitive to the level of disorder in auditory and visual sequences, demonstrating higher activity the more ordered the sequence^50^. Orbitofrontal cortex is proposed to be a source of top-down modulation on auditory cortex according to context^51^.

From 165-245 ms, the increased response to deviants in REG was associated with greater left temporal and right prefrontal activity. Left middle temporal gyrus is sensitive to unexpected uncertainty, a quantity which reflects the strength of expectation violation given knowledge of stimulus reliability, during a decision-making task^52^, although the authors found a reduction in activity with surprise. Right prefrontal activity is associated with signalling prediction violation in a variety of domains^17,53,54^, and was recently proposed as a locus of integration of confidence and prediction violation^55^.

Overall, the results replicate the ubiquitous network of bilateral auditory cortex and right pre-frontal sources as underpinning frequency-based deviance detection and additionally implicate the temporal pole as well as right orbitofrontal and pre-frontal cortex in nuancing these responses according to the preceding sequence context.

In addition to the effect of regularity on deviant processing, we also observed an overall larger sustained response to REG relative to RAND patterns. This stands in contrast to the standard observation of a reduction in brain responses to predictable sounds^16,18,21,30,38^. Because in most of those cases, ‘predictability’ was implemented using single repeated tone-pips (e.g. ‘standards’ in the oddball paradigms), it is likely that adaptation, at least partially, contributes to this effect1^12,21^; but see Todorovic & de Lange^56^). In contrast, the present paradigm uses rapid, wide-band, regularly-repeating sequences which decouple the effects of repetition from those associated with predictability per se and our results are consistent with previous reports using similar stimuli^27,28,57^.

A specific mechanistic account for the increased sustained response remains elusive, but previous work has demonstrated that the amplitude of the sustained response is related to the predictability or precision of the ongoing acoustic pattern^27,28,57,58^, such that increased predictability is systematically associated with higher sustained responses. This effect, underpinned by increased activity in a network of temporal, frontal and hippocampal sources^27,58^, may reflect a mechanism which tracks the context-dependent reliability of sensory streams.

According to predictive coding theory, surprise is determined by two processes: *prediction error* evoked by a stimulus that differs from expectations, and also the *precision* associated with the input; i.e. the reliability attributed to the sensory stream^14,59^. It is hypothesized that brain responses to predictable (highly precise) stimuli are up-weighted (e.g. through gain modulation) to focus perception on stable features of the environment^60^. It is tempting to interpret the increased amplitude of the sustained response to regular sequences as a manifestation of precision-weighting^27,28,57,58^, though it remains unclear whether the sustained effects seen here are indeed excitatory (as the gain modulation postulated by predictive coding). Alternatively, the observed effects may reflect increased *inhibition* (see also discussion in Southwell et al, 2017; Barascud et al, 2016), consistent with emerging findings which implicate inhibitory interneurons in controlling the coding of predictable sound sequences^61,62^.

Importantly, the increased deviant response during regular sequences was observed in addition to an increase in the ongoing sustained response. Furthermore, source analysis suggests the response to deviants in regular sequences was not merely enhanced relative to matched deviants in random sequences but rather arose in part via the involvement of distinct underlying sources. Therefore, an account in terms of differential precision weighing over the same prediction error units, as proposed by predictive coding^59,60^, may not fully account for the observed effects. Instead, the results are consistent with a predominantly auditory cortical source coding for frequency deviance and more frontal sources associated with encoding more abstract properties of pattern violation.

## Methods

### Stimuli

Stimuli consisted of 50-ms tone pips of varying frequency, arranged in regular (REG) or random (RAND) frequency patterns over a total duration of 3000ms. Frequencies were drawn from a pool of 26 logarithmically-spaced values between 198 and 3563Hz (12% increase in frequency at each step; equivalent to two musical semitones). To generate each sequence, 13 adjacent frequencies were chosen at random from the larger pool (see Figure 1a) and then a random subset of 10 of these frequencies were retained, so that all sequences had a similar bandwidth and contained exactly 10 unique frequencies (‘alphabet size’ = 10). REG sequences were generated by permuting the 10 chosen frequencies and then repeating that order six times (Figure 1b; upper left). Matched RAND sequences were generated by shuffling each REG sequence, with the constraint that no two adjacent tones were the same frequency (Figure 1b; upper right). Overall, the stimulus generation procedure ensured that REG and RAND sequences are matched exactly in terms of the first order distribution of tones; the only difference being whether they are arranged in a predictable (REG) or unpredictable (RAND) order.

Half of the sequences (henceforth denoted as REG_dev_ and RAND_dev_) contained a single ‘deviant’ tone between 1500 and 2750ms post-onset (chosen at random for each stimulus), which is equivalent to a minimum of 3 REG cycles (Figure 1b; lower panels). Our previous work^27,28^ determined that the detection of regularity and the associated brain responses take place between 1-2 cycles. A latency of 3 cycles therefore assures that the processing of the regular pattern has stabilized (see also Figure 2c). The deviant tones replaced the corresponding standard tone. The deviant frequency was either higher or lower than the range spanned by the 10 ‘standard’ frequencies in the sequence, with a minimum distance of two frequency steps. Throughout the entire set of stimuli all 26 frequencies could be deviants or standards. Furthermore, to ensure all ten standard frequencies were approximately equally probable before the deviant, RAND_dev_ were generated by shuffling separately before and after the chosen deviant position. Stimuli were generated in matched sets of four (REG, RAND, REG_dev_, RAND_dev_), using the same ‘alphabet’ for standards (and the same frequency for the deviants, if applicable). Sequences were unique on each trial and generated anew for each subject.

### Experiment 1 – behavioural sensitivity to deviants in REG and RAND sequences

#### Stimuli and Procedure

Subjects heard 96 trials each of REG, RAND, REG_dev_ and RAND_dev_ (in random order), and were instructed to respond by button press when they heard a deviant tone. Fourty-eight additional control trials were also included, with the same number and timing of tone pips, but consisting of a single, repeating standard frequency (CTRL). Twenty-four of these contained a deviant tone at least 2 whole tones away from the standard (CTRL_dev_); deviant and standard frequencies were chosen at random for each stimulus. Subjects were instructed to respond by button press as quickly as possible when a deviant tone was detected. Trials were presented in a random order, but the proportion of each condition across each block of 72 trials was kept the same. The testing session was preceded by a practice session of 28 trials; conditions were the same as the main experiment and in the same proportions.

#### Analysis

Dependent measures are d’ scores^63^ and response times (RT; measured between the onset time of the deviant and the subject’s key press). Outlier trials deviating from the condition-wise mean reaction time by more than 2 standard deviations were excluded; this resulted in exclusion of no more than 6% of trials for each condition. Sensitivity scores (d’) to deviants in each condition were calculated using the hit and false alarm rates. In cases where either rate was 0 or 1, a half trial was (respectively) added or subtracted to the numerator and denominator of the rate calculation; to avoid infinite d’ values.

#### Participants

10 paid participants took part (age 18-34, mean 24.4 years; 5 female). All (here and in Experiment 2 below) were right handed and reported normal hearing and no history of neurological disorders. The experimental protocol for all reported experiments was approved by the University College London research ethics committee.

### Experiment 2 – EEG responses to deviants within REG and RAND contexts

#### Stimuli and procedure

The stimulus set comprised four sequence types: REG, RAND, REG_dev_ and RAND_dev_, as described above. These were presented to naïve, distracted listeners whilst their brain activity was recorded with EEG. As in Experiment 1, each trial was unique and sequences were generated anew for each subject. A total of 600 sequences were presented; 150 of each condition. The session was split into 6 blocks to provide breaks, each with 25 trials per condition presented in a random order. The inter-trial interval (ISI) was jittered between 1100 and 1500 ms. Stimuli were presented with the Psychophysics Toolbox extension in Matlab^64^, using EarTone in-ear earphones with the volume set at a comfortable listening level. During the experiment, subjects watched a subtitled film of their choice, with the audio muted. They were informed that there would be some sounds played during the session, and were presented with a single example of RAND as a demonstration, but were instructed to ignore all sounds.

Following the session, subjects were asked the following questions about the sounds they heard:

1. During the EEG experiment, you heard some sounds. How distracting did you find them (1 = not at all, 5 = very distracting all the time)
2. Please describe the sounds briefly – what did you notice?
3. Did you hear any patterns in the sounds?
4. Did you hear any beeps that broke the pattern?

#### EEG Recording and Analysis

EEG was recorded using a 128-electrode Biosemi system (Biosemi Active Two AD-box ADC-17, Biosemi, Netherlands) at a sampling rate of 2048 Hz. Data were pre-processed and analysed using Fieldtrip (http://www.fieldtriptoolbox.org/)^65^ toolbox for MATLAB (2015a, MathWorks). Separate analysis pipelines were used to analyse the whole sequence response (time-locked to sequence onset) and the deviant response (time-locked to the onset of the deviant tone). All filtering was performed with a two-pass Butterworth filter.

For the whole sequence analysis, data were high-pass filtered at 0.1Hz (third-order) and divided into 5000-ms epochs (with 1000 ms pre-stimulus-onset and 1000 ms post-offset). Subsequently, all data were low-pass filtered at 100 Hz (fifth-order), down-sampled at 200 Hz, and baseline-corrected relative to the pre-onset interval. Outlier epochs were removed on the basis of summary statistics (variance, range, maximum absolute value, z-score, maximum z-score, kurtosis) using Fieldtrip’s visual artefact rejection tool. On average 5% of epochs were removed for each subject (range 0-10%). Artefacts related to eye movements, blinks and heartbeat were identified using independent component analysis (ICA) on a copy of the data that had been high-pass filtered at 1Hz before epoching (optimal for identifying artefacts with ICA^66^), but the expected slow dynamics of the sequence-evoked response is only visible with a lower cutoff of e.g. 0.1 Hz^27,28^. Any channels previously identified as noisy were not included in the ICA procedure. The obtained unmixing matrix was then used to estimate component timeseries in the 0.1-Hz filtered data. The artefactual components were removed for each subject to yield the cleaned dataset. Missing bad channels were reconstructed as the average of their immediate neighbours. Subsequently the data were re-referenced to the mean of all channels, averaged over epochs of the same condition, baseline-corrected (200 ms preceding stimulus onset) and low-pass filtered at 30Hz (fifth-order) for plotting and analysis.

For quantifying the deviant response, data were high-pass filtered at 2Hz (third-order) and divided into 2000-ms epochs, with 1000 ms on either side of the deviant onset time.

Conditions without a deviant (REG and RAND) were epoched relative to the average deviant timing; rounded down to the nearest tone onset, i.e. 2100ms. Subsequent analysis was identical to the one described for the whole sequence analysis (above).

For the offset response analysis, the sequence-evoked data were high-pass filtered at 2Hz, re-aligned into epochs (2800-3500 ms) and baseline-corrected based on the interval 2800-3000 ms.

#### Statistical analysis

To assess the effect of deviance, first REG and RAND conditions were pooled; splitting the data into deviant (REG_dev_ and RAND_dev_) and no-deviant (REG and RAND) epochs. A two-tailed t-test was used to quantify the effect of deviance (deviant minus no-deviant epochs) in each time point and each channel. We used Fieldtrip’s cluster-based permutation test (using a two-tailed dependent-samples t-test with p=0.025), which takes spatial and temporal adjacency into account, to correct for multiple comparisons^65,67^. The significance threshold was chosen to control family-wise error-rate (FWER) at 5%.

To characterize the overall sequence-evoked response to REG and RAND, the root mean square (RMS) of the evoked potential over all channels was calculated for each time point to give a time-series which reflects the instantaneous power of the evoked response^27,28^. The distribution of RMS across subjects (mean, standard error) was then estimated for each condition using bootstrap resampling across subjects^68^ with 1000 iterations, for plotting of the group average response in Figure 3a. The significance of the difference in RMS between REG and RAND was assessed using the same cluster-based permutation statistics as for the deviant response, at each time sample, from sequence onset to 500ms following offset. T-tests (2-tail) were performed using t-statistics computed on clusters in time, and controlled for a family-wise error rate of 0.05^67^.

#### Source Analysis

To visualise estimated source activity during the observed deviant effects, source analysis was performed to generate the source-level T-maps shown in Figure 2; In the absence of individual structural scans, a head model derived from a template MNI brain was used (colin27; as included in the Fieldtrip toolbox) for which the volume conductance model was computed from MRI images using the Boundary Element Method^69^. A triangulated cortical sheet, with 5124 vertices, derived from this scan was used as the source model. Source inversion was performed on individual subjects and separately for each condition, using Minimum Norm Estimation (MNE)^70^. The analysis focused on 0-300ms relative to the onset of the deviant. Source data were averaged within the time intervals corresponding to the two windows identified in sensor space (80-145 and 165-245 ms respectively). Subsequently, T-statistic maps were computed for the main effect of deviance (deviant > no-deviant), and the orthogonal contrast of regularity i.e. (REG_dev_ – REG) > (RAND_dev_ – RAND), within each time window. Data were interpolated onto an inflated cortical surface for visualisation in Figure 2e & 2f and are presented using a threshold of T=2. No statistical analysis was carried out on the thresholded source data; as the time-windows were defined by statistically significant effects in sensor space. Due to the limited precision afforded by the template-based source modelling used here, we discuss activation patterns in terms of general areas as opposed to specific MNI coordinates.

#### Participants

Data from 20 paid subjects are reported (age 19-32, mean 22.8 years. 9 female). None had participated in the behavioural task (Experiment 1). One additional subject was excluded from analysis due to excessively noisy data.

